# Integrity and stability of virulence plasmids from *Pseudomonas syringae* are modulated by mobile genetic elements and multiple toxin-antitoxin systems

**DOI:** 10.1101/453399

**Authors:** Leire Bardaji, Maite Añorga, Myriam Echeverría, Cayo Ramos, Jesús Murillo

**Author notes:** Corresponding author: Jesús Murillo.

## Abstract

**Background:** Virulence plasmids are critically exposed to genetic decay and loss, particularly in *Pseudomonas syringae* strains because of their high content of mobile genetic elements and their exploitation of environmental niches outside of the plant host. The demonstrated high plasticity and adaptability of P. syringae plasmids, involving the acquisition and loss of large DNA regions, contrasts with their usual high stability and the maintenance of key virulence genes in free living conditions. The identification of plasmid stability determinants and mechanisms will help to understand their evolution and adaptability to agroecosystems as well as to develop more efficient control measures.

**Results:** We show that the three virulence plasmids of *P. syringae* pv. savastanoi NCPPB 3335 contain diverse functional stability determinants, including three toxin-antitoxin systems (TA) in both pPsv48A and pPsv48C, whereas one of the two replicons of pPsv48C can confer stable inheritance by itself. Loss of pPsv48A increased by two orders of magnitude upon functional inactivation of its TA systems. However, inactivation of the TA systems from pPsv48C did not result in its curing but led to the recovery of diverse deletion derivatives. One type consisted in the deletion of an 8.3 kb fragment, with a frequency of 3.8 ± 0.3 × 10^−3^, by recombination between two copies of MITE*Psy2*. Likewise, IS*801* promoted the occurrence of deletions of variable size by one-ended transposition with a frequency of 5.5 ± 2.1 × 10^−4^, 80 % of which resulted in the loss of virulence gene *idi*. These deletion derivatives were stably maintained in the population by replication mediated by *repJ*, which is adjacent to IS*801*. IS*801* also promoted deletions in plasmid pPsv48A, either by recombination or one-ended transposition. In all cases, functional TA systems contributed significantly to reduce the occurrence of plasmid deletions *in vivo*.

**Conclusions:** Virulence plasmids from *P. syringae* harbour a diverse array of stability determinants with a variable contribution to plasmid persistence. Additionally, multiple TA systems favour the long-term survival and integrity of virulence plasmids, as well as the maintenance of pathogenicity genes in free-living conditions. This strategy is likely widespread amongst native plasmids of *P. syringae* and other bacteria.

## Background

Plasmids are dispensable extrachromosomal elements widely distributed in bacteria, facilitating their survival and the colonization of eukaryotic hosts [1–4]. The plasticity and transmissibility of plasmids contribute to a rapid dissemination of resistance and virulence genes, thus promoting the emergence of uncontrollable bacterial diseases, both in clinical and agricultural settings [5–9]. However, plasmids are usually large and exist in several copies per cell, potentially imposing a significant metabolic burden to the cell, which might facilitate the emergence of plasmid-free derivatives in the absence of selection for plasmid-borne characters [8, 10, 11]. This metabolic cost can be lowered by diverse plasmid-host adaptations, such as deletions, mutations in the plasmid replication machinery, or chromosomal mutations [8, 10, 11]. Additionally, plasmids can increase their stability by conjugal transfer and/or by carrying a battery of specifically dedicated genetic determinants, classified into three main categories [12–14]. Partition determinants, in the first category, direct the active segregation of plasmid molecules during cell division. All low-copy plasmids appear to contain a partition system, which usually consists of an operon of two genes plus a specific DNA sequence for recognition. Multimer resolution systems comprise the second category and include recombinases that resolve plasmid cointegrates and maximize the number of plasmid copies available at cell division. The third category, postsegregational killing systems, include toxin-antitoxin (TA) systems and, less prominently, restriction modification loci; these systems ensure plasmid maintenance by inhibiting cell growth.

The *Pseudomonas syringae* complex is considered the most important bacterial plant pathogen in the world [15]. Most strains contain plasmids with an array of adaptive genes that increase aggressiveness, expand their host range, and confer resistance to antibacterials or to UV light [3, 7, 16–18]. Most of these plasmids belong to the so-called PFP group, characterized by sharing the highly conserved RepA-PFP replicon. These replicons are highly plastic and adaptable, and strains often contain two or more stably co-existing PFP plasmids [7, 19–21]. Insertion sequences, transposons and miniature inverted-repeat transposable elements (MITEs) can account for up to at least a third of a PFP plasmid, actively participating in the acquisition and exchange of adaptive characters [20–24]. Insertion sequence IS*801* (1.5 kb), and its isoforms, is particularly significant because of its relatively high transposition frequency, its common association with virulence genes and its ability to undergo one-ended transposition, whereby the element can mobilize adjacent DNA [22, 24, 25]. Additionally, plasmids of *P. syringae* have a mosaic structure and often share extensive regions of similarity, suggesting their evolution through the acquisition and loss of large DNA regions in a multistep process [17–20, 23, 26]. Despite this, plasmid profiles of individual strains appear to be characteristic and stable, although certain plasmids can be lost with high frequency under certain culture conditions [3, 27–30]. In fact, diverse potential stability determinants were identified among PFP plasmids [18, 20, 21, 31–33], although it is not yet clear whether or not they are functional and what their role in the bacterial life cycle is.

*P. syringae* pv. savastanoi NCPPB 3335 causes tumours in olive (*Olea europaea*) and is a prominent model for the study of the molecular basis of pathogenicity on woody hosts [34, 35]. This strain contains three PFP virulence plasmids pPsv48A (80 kb), pPsv48B (45 kb) and pPsv48C (42 kb) [21]. Plasmid pPsv48A carries the virulence gene *ptz*, involved in the biosynthesis of cytokinins, and the Type III effector gene *hopAF1*; pPsv48B carries the Type III effector gene *hopAO1* and, in turn, plasmid pPsv48C carries the virulence gene *idi*, potentially involved in cytokinin biosynthesis. Both pPsv48A and pPsv48C are essential for the production of tumours in olive plants [21, 36], whereas pPsv48B contributes to fitness and virulence *in planta* [37]. Although pPsv48A and pPsv48B can be cured, pPsv48C is remarkably stable and could not be evicted from strain NCPPB 3335 [21], perhaps because it carries two different replicons [38]. We were interested in the identification and characterization of the stability determinants of the plasmid complement of strain NCPPB 3335, to gain insights into the mechanisms allowing the coexistence of PFP plasmids and the dynamics of virulence genes.

Here, we determined that the stability of plasmids pPsv48A and pPsv48C is modulated by diverse mechanisms. The presence of two replicons in pPsv48C contributes to its very high stability, whereas three TA systems in both plasmids A and C are necessary to avoid the occurrence of high-frequency deletions and rearrangements mediated by an IS*801* isoform and MITE*Psy2*, while simultaneously favouring the maintenance of virulence genes *ptz* and *idi* when outside the plant.

## Results

### Identification of functional stability determinants in the three native plasmids

We selected 28 coding sequences from pPsv48A, pPsv48B and pPsv48C that could be involved in plasmid maintenance (Table 1). These were organized in eight putative TA systems (operons TA1 to TA8) and seven other potential stability determinants (SD1 to SD7). All these were cloned into pKMAG-C either individually or in groups, when suspected to form an operon, and the stability conferred to the vector was assayed in *P. syringae* pv. syringae B728a (Fig. 1). pKMAG-C is able to replicate in both *E. coli* and pseudomonads [38], and is highly unstable in *P. syringae*.

**Table 1.**
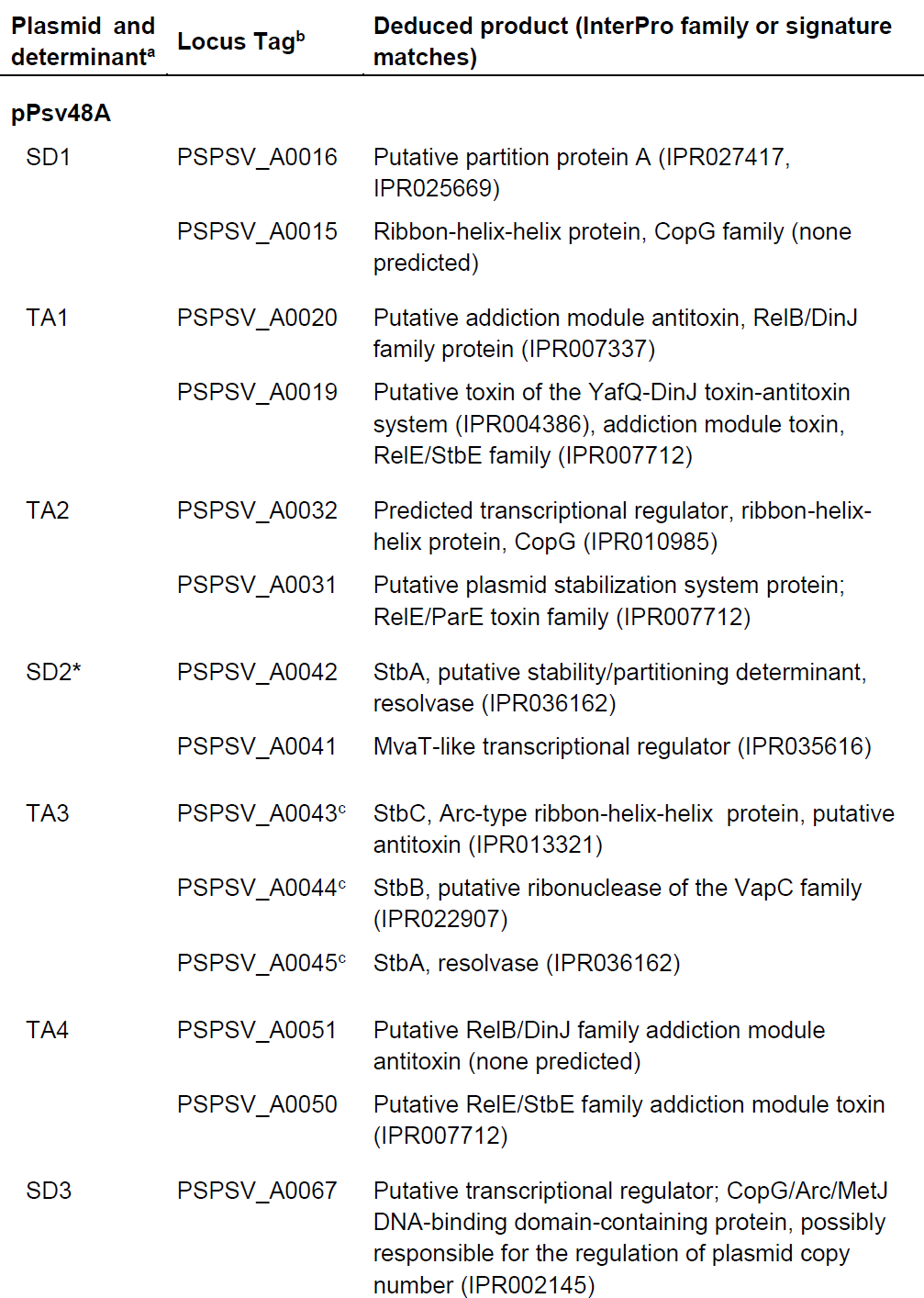
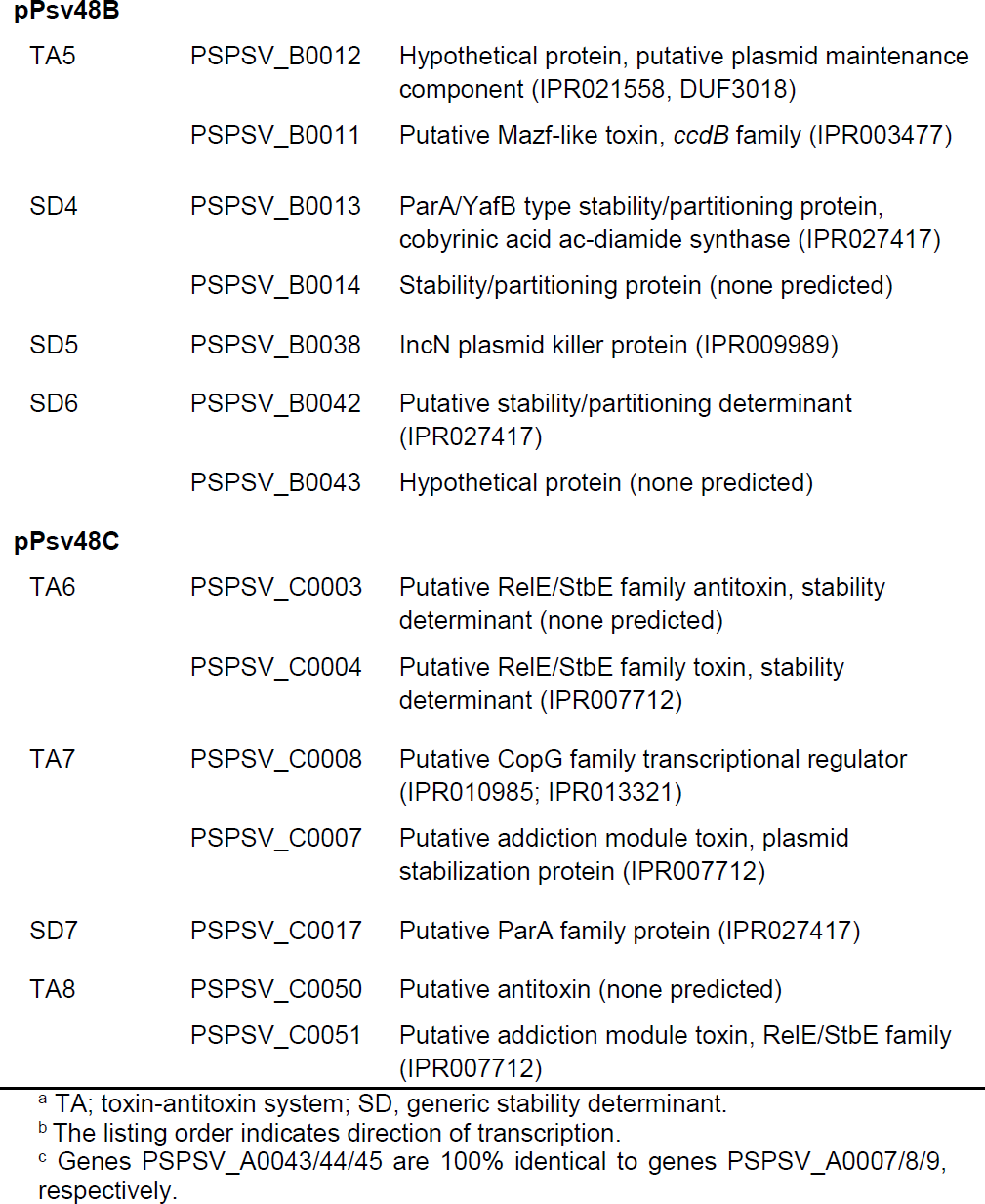
Putative stability determinants identified in the three native plasmids of *P. syringae* pv. savastanoi NCPPB 3335.

**Fig. 1.**
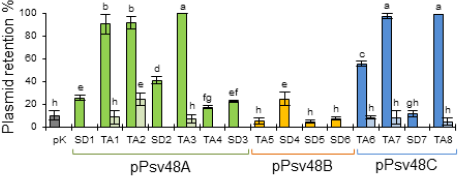
Functional analysis of putative stability determinants. Bars indicate the average percentage of *P. syringae* pv. syringae B728a cells retaining pKMAG-C (pK) or the indicated putative stability determinants from the three native plasmids of *P. syringae* pv. savastanoi NCPPB 3335 (Table 1) cloned into pKMAG-C. For TA systems leading to >50 % of plasmid retention, we show to their right the stability conferred by the corresponding antitoxin cloned in pKMAG-C. Bars topped with different letters indicate means that are significantly different according to a one-way ANOVA followed by Duncan’s multiple range test (P < 0.05).

All seven determinants tested from pPsv48A (Table 1), significantly increased stability of pKMAG-C to varying degrees (Fig. 1). Three are TA systems and, as expected, conferred very high levels of stability only when cloned completely, but not when the putative antitoxin was cloned by itself (Fig. 1), although the antitoxin from system TA2 on its own conferred moderate levels of stability. System TA3 is widespread in pseudomonads [e.g., 33, 39, 40] and it is an operon of the TA genes *stbCB* plus the putative resolvase *stbA* (Table 1). Constructs containing either *stbCBA* or only genes *stbCB* conferred equal high levels of stability (not shown); therefore, we evaluated the possible contribution of *stbA* to stability by cloning it separately. *stbA* is the last CDS in the *stbCBA* operon and predictably lacks a promoter; thus, we tested functionality of the *stbA* allele PSPPH_A0042, which is the first CDS of another putative operon (SD2 in Fig. 1) and shows 90 % nt identity to the allele in operon *stbCBA*. Operon SD2 also significantly increased stability of pKMAG, likely through resolution of plasmid multimers by the StbA resolvase [13], suggesting that operon *stbCBA* might contribute to stability through different mechanisms.

Only one of the four determinants from pPsv48B evaluated here (Table 1) appeared to contribute, albeit modestly, to plasmid stability (Fig. 1). This was unexpected because low-copy number plasmids usually carry diverse maintenance determinants [41]. Nevertheless, it is possible that this plasmid contains stability genes that were not included or whose activity was not detected in our assays, and/or that its stability is increased by conjugation [12].

Three TA systems, out of the four determinants tested from pPsv48C, contributed to plasmid stability (Table 1); again, the putative antitoxins did not confer any stability by themselves (Fig. 1). Remarkably, the different TA systems showed distinct behaviours in our assays, which varied from no apparent contribution to stability (TA5; Fig. 1) to conferring moderate to very high stability levels (e.g. TA4 and TA3 or TA8, Fig. 1).

### The two replicons from pPsv48C confer distinct stability

To explore the basis of the very high stability of pPsv48C, we evaluated the contribution of the RepA-PFP and RepJ replicons to its maintenance. Therefore, we cloned them into the *E. coli* vector pKMAG and evaluated stability in the plasmidless strains *P. syringae* pv. syringae B728a and *P. syringae* pv. savastanoi UPN912 (Fig. 2). Strain UPN912 derives from NCPPB 3335 by curing of its three native plasmids (Table 2).

**Fig. 2.**
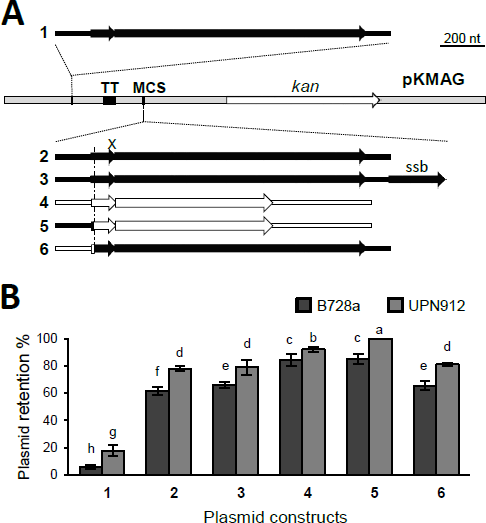
Stability of constructs containing the native RepA-PFP and RepJ replicons from pPsv48C, and their chimeras. (A) Fragments of the RepA-PFP (black) or RepJ (white) replicons, and their chimeras, were cloned at the indicated positions into pKMAG; small and large arrows represent the putative leader peptide and the replication initiator genes, respectively. TT, T4 transcription terminator; MCS, multiple cloning site; *kan*, kanamycin resistance gene. (B) Bars indicate the average percentage of *P. syringae* pv. syringae B728a cells (dark grey) or of *P. syringae* pv. savastanoi UPN912 cells (light grey) retaining each construct. Numbers indicate the constructs shown in panel A. Bars topped with different letters indicate means that are significantly different according to a two-way ANOVA followed by Duncan’s multiple range test (P < 0.05).

**Table 2.**
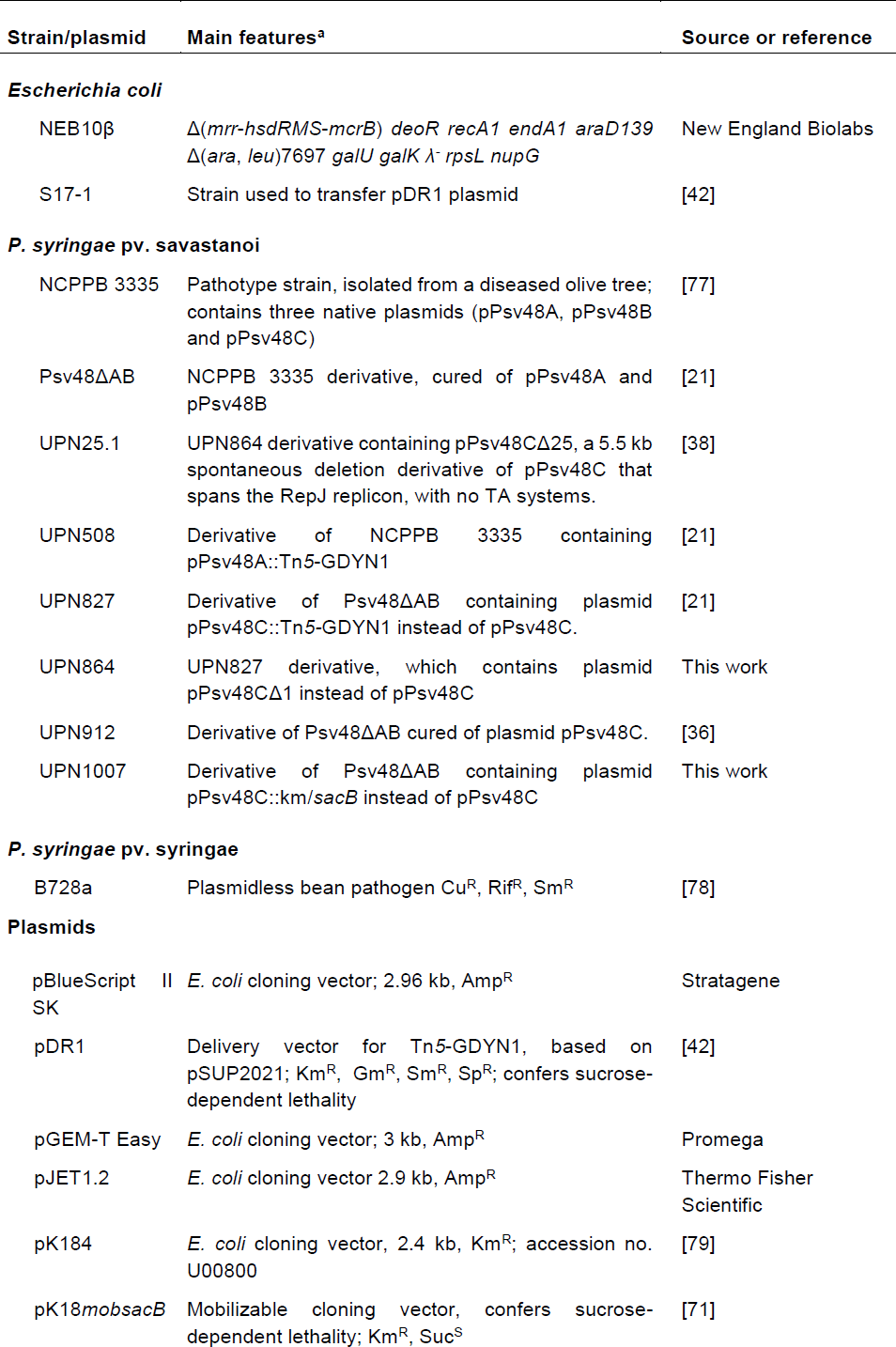
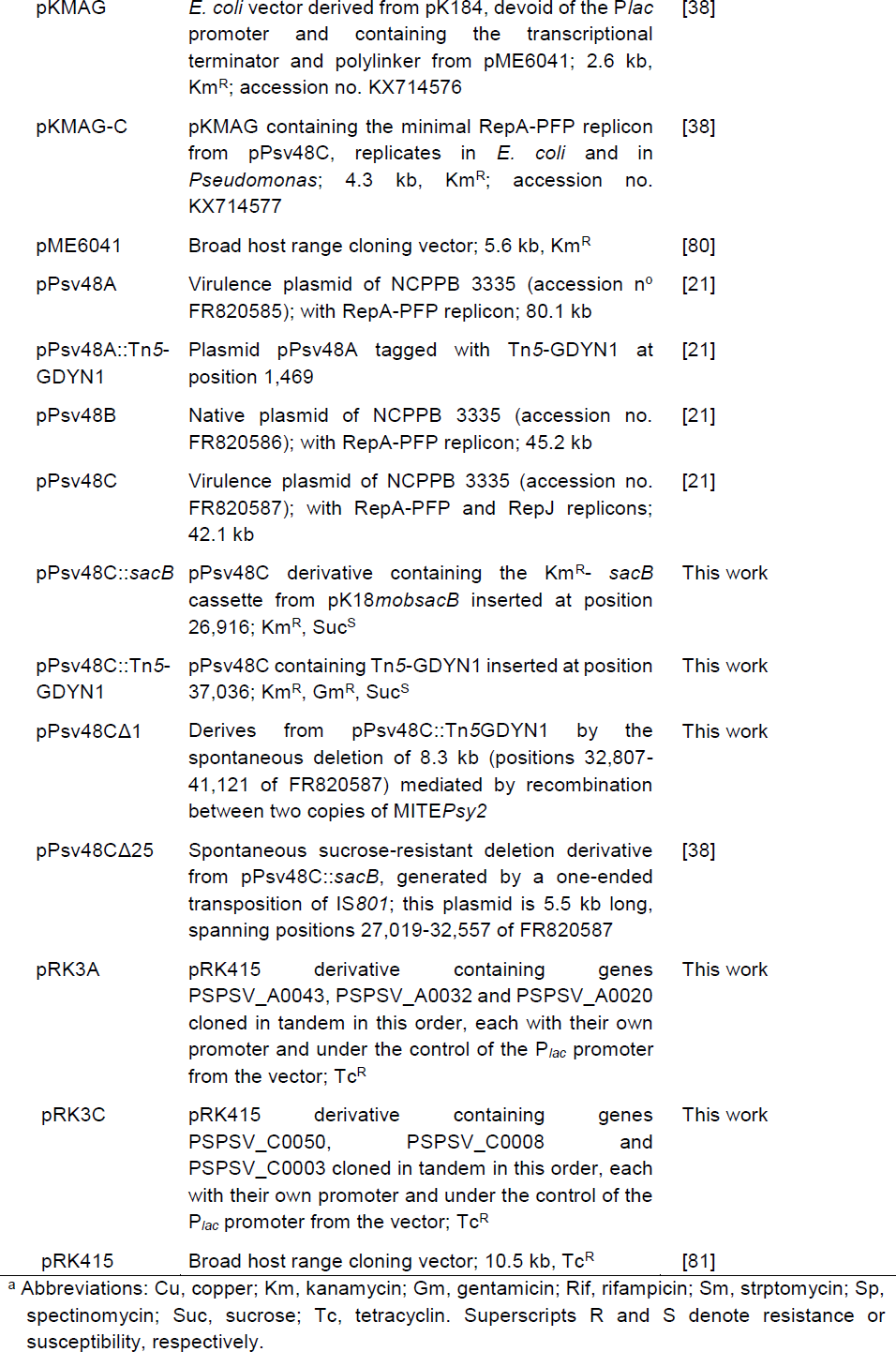
Bacterial strains and plasmids used in this study

Construct pKMAG-C, containing the RepA-PFP replicon cloned outside the polylinker of the vector, was highly unstable and was nearly completely lost after only one night of growth (Figs. 1 and 2). However, its cloning after the transcription terminator of pKMAG significantly increased stability (2 in Fig. 2). Gene *ssb*, which is frequently found downstream of the *repA* gene [20, 21, 32] only showed a marginal contribution to stability (compare 2 and 3, Fig. 2). In turn, the RepJ replicon conferred a significantly higher stability than the RepA-PFP replicon (compare 2 and 4, Fig. 2). Noticeably, all the constructs were significantly more stable in strain UPN912 than in B728a (Fig. 2), suggesting that these replicons are adapted to the bacterial host in which they occur naturally, to maximize their survival.

The RepA-PFP and RepJ replicons consist of two separable functional fragments: a control region, containing the promoter, a putative antisense RNA and a leader peptide, and a replication region, containing the replication initiator protein (*rep*) gene [38]. The approx. 0.3 kb control region determines the transcription rate of the *rep* gene. The RepA-PFP and RepJ replicons share very similar, but not identical control regions preceding the *rep* gene [38], and we hypothesized that this could potentially influence replicon stability. We therefore evaluated the stability of constructs containing chimeric replicons, with the replication control region (Rex-C module) reciprocally swapped [38]. The highest stability in UPN912, but not in strain B728a, was reached with the chimera RepA-PFP:RepJ (control:replication modules; construct 5, Fig. 2), indicating that replicon stability is mostly dependent on the activity of the replication module, but it can be modulated by the control module (Fig. 2).

The significant values of plasmid loss observed for RepJ (Fig. 2) conflicted with the high stability observed for pPsv48C deletion derivatives (not shown), suggesting that we did not clone all the replicon sequences needed for stable replication. We therefore tested the stability of a spontaneous 5.5 kb deletion derivative of pPsv48C (clone pPsv48Δ25; Table 2), containing the minimal RepJ replicon [38] plus additional DNA that did not include any other potential plasmid maintenance genes. Plasmid pPsv48Δ25 was maintained in 100% of the cells obtained from starting cultures and after seven sequential culture transfers (1622 and 2804 colonies tested, respectively). In contrast, the RepJ construct in pKMAG (construct 4 in Fig. 2) was retained by 94 ± 2 % of UPN912 cells from starting cultures and by only 63 ± 2 % of the cells after seven transfers (2366 and 2666 colonies tested, respectively). These results indicate that the native RepJ replicon is larger than the minimal replicon [38] and underscore its high stability in its genetic context.

### A toxin-antitoxin system prevents a deletion in pPsv48C mediated by MITEs

We sought to obtain derivatives of NCPPB 3335 cured of plasmid pPsv48C, and to evaluate the contribution of its three TA systems to stability. We thus constructed strain UPN827, containing a transposon carrying the *sacB* gene (Tn*5*-GDYN1) inserted into pPsv48C (Fig. 3; Table 2); this allowed us to easily select for plasmid loss by growth in the presence of sucrose [42]. To inactivate functionally the TA systems [43] and facilitate plasmid loss, we constructed pRK3C, containing the three antitoxin genes from pPsv48C cloned in pRK415 (Table 2), and introduced it into UPN827 to neutralise the three corresponding toxins.

**Fig. 3.**
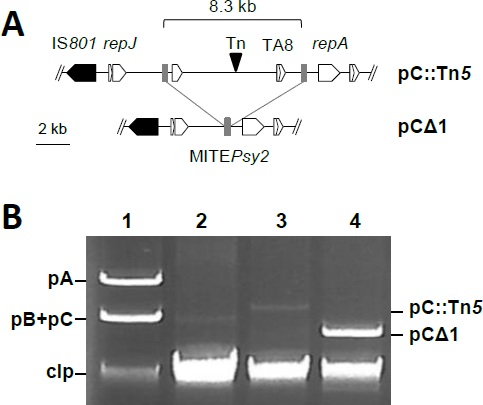
Recombination between two directly repeated copies of MITE*Psy2* causes a deletion on pPsv48C. (A) Partial map of pPsv48C::Tn*5*-GDYN1 (pC::Tn*5*) showing the relative positions of its only copy of the IS*801* isoform, its two replication initiation protein genes (*repJ* and *repA*), and toxin-antitoxin system 8 (TA8). Small grey rectangles, MITE*Psy2*; inverted black triangle, Tn*5*-GDYN1 (Tn). pCΔ1 is pPsv48CΔ1, containing an 8.3 kb deletion resulting from MITE*Psy2* recombination. (B) Electrophoresed uncut plasmid preparations from: (1) *P. syringae* pv. savastanoi NCPPB 3335; (2) Psv48ΔAB; (3) UPN827, and (4) UPN864. pA, pPsv48A; pB; pPsv48B; pC, pPsv48C; pCΔ1, pPsv48CΔ1; clp, chromosomal DNA and linearized plasmids. Lanes were loaded with equivalent amounts of cell lysates; results are representative of at least 20 independent plasmid preparations.

We routinely obtained 50 times more sucrose-resistant colonies with strain UPN827(pRK3C) (38 ± 3 × 10^−4^ suc^R^ colonies) than with its parental strain UPN827(pRK415) (0.8 ± 0.4 × 10^−4^ suc^R^ colonies), and this difference was significant. All suc^R^ colonies examined contained an 8.3 kb deletion in pPsv48C caused by the recombination of two direct copies of MITE*Psy2,* as assessed by sequencing, eliminating the *sacB* transposon Tn*5*-GDYN1 (Fig. 3A). One of these plasmids was retained and designated pPsv48CΔ1 (Fig. 3A). These results indicate that, despite its small size (228 nt), MITE*Psy2* is a hot spot for recombination.

In plasmid profile gels of the wild type strain NCPPB 3335, pPsv48C routinely appears with lower intensity than the other two native plasmid bands [21] (Fig. 3B). Remarkably, bands of plasmid pPsv48CΔ1 were repetitively more intense than those of the wild type plasmid or of pPsv48C::Tn*5*-GDYN1 (Fig. 3A), suggesting that the 8.3 kb deletion caused a higher copy number. We estimated a moderate copy number for plasmids pPsv48A (8.0 ± 1.0), pPsv48B (8.6 ± 1.6) and pPsv48C (6.6 ± 1.2), with no significant differences among them. These are as expected for medium-size native plasmids [44] and similar to the five copies reported for the native plasmid pFKN from *P. syringae* pv. maculicola [23]. Unexpectedly, the estimated copy number of pPsv48CΔ1 (6.9 ± 0.8) was not significantly different from that of pPsv48C. These results indicate that each of the three native plasmids from strain NCPPB 3335 exist in 6-9 copies per cell, and that the 8.3 kb fragment from pPsv48C does not carry any determinant involved in copy number control. This also suggests that structural differences among plasmids could differentially impact their purification by alkaline lysis and questions the use of agarose gel electrophoresis to estimate relative plasmid DNA quantities.

### Toxin-antitoxin systems from pP sv48C prevent accumulation of plasmid deletions mediated by IS*801*

Our preliminary experiments soon indicated that the inactivation of the three TA systems of pPsv48C did not facilitate the isolation of plasmid-cured strains but, instead, led to the recovery of deletion derivatives generated by one-ended transposition of the IS*801* isoform CRR1 (Fig. 4) [21]; for clarity, we will henceforth refer to this isoform as IS*801*. Therefore, strain UPN1007 was used to better estimate the causes and frequency of the different deletions. This strain carries plasmid pPsv48C::*sacB*, containing a Km^R^-*sacB* cassette immediately adjacent to the only IS*801* copy of pPsv48C (Fig. 5); thus, the selection of suc^R^ colonies would allow for the identification and quantification of all types of deletions mediated by one-ended transposition of IS*801*.

**Fig. 4.**
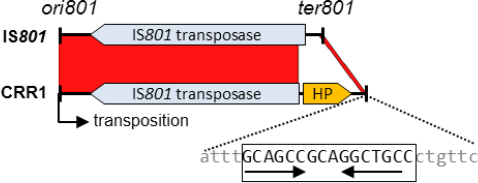
Comparison of the wild type IS*801* with its isoform CRR1. Blastn alignment of IS*801* (X57269; 1512 nt) and CRR1 (from FR820587; 1765 nt); the red bands connecting the two elements indicate collinear regions of identify. CRR1 contains an insertion of 365 nt, causing a deletion of 112 nt that removes the predicted transposase start codon and leaves 26 nt in the *ter801* terminus (expanded sequence). This 26 nt region contains a conseverd motif (captial letters) with an inverted repeat sequence (horizontal arrows), probably involved in recognition and interaction with the transposase [45].

**Fig. 5.**
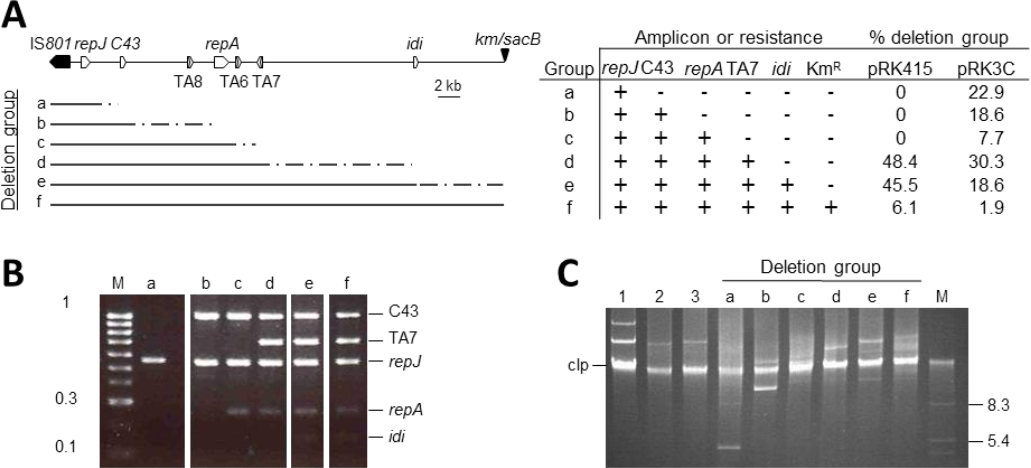
Types of deletions of pPsv48C::*sacB* as influenced by functional toxin-antitoxin systems. (A) Left: Map of pPsv48C::*sacB*; TA6, TA7 and TA8, toxin-antitoxin systems; C43, locus PSPSV_C0043; inverted triangle, Km^R^-*sacB* cassette cloned 0.1 kb 3’ of the IS*801* isoform. Lines under the map indicate the minimum (black line) and maximum (dotted line) extent of DNA transported by IS*801* on each group of suc^R^ plasmids. Right: Presence (+) or absence (−) of specific amplicons for each of the genes shown, or of resistance (+) and sensitivity (−) to kanamycin. Last two columns indicate the percentage of suc^R^ colonies containing each plasmid group in UPN1007 containing pRK3C, leading to functional inactivation of the TA systems, or the empty, or the empty vecor pRK415. Gel showing typical patterns if multiplex PCRamplifications (B) and uncut plasmids (C) of example clones from each plasmid group. M, molecular weight markers, in kb; clp, chromosomal DNA and linearized plasmids. Lanes: (1) *P. syringae* pv. savastanoi NCPPB 3335; (2) Psv48ΔAB, containing only pPsv48C, and (3) UPN864, containing only pPsv48C::*sacB*

The frequency of sScuc^R^ colonies was 1.8 ± 0.7 × 10^−4^ for UPN1007 containing the empty vector but significantly higher (5.5 ± 2.1 × 10^−4^) for strain UPN1007(pRK3C), in which the three TA systems are functionally inactivated (Fig. 5). The plasmid profile and PCR analyses of >700 independent clones, plus sequencing of 13 of them, indicated that none had lost pPsv48C but showed a plasmid band of *ca*. 4 to 42 kb resulting from deletions of variable size in this plasmid. All deletion derivatives contained IS*801* and *repJ* (Fig. 5), and sequencing showed that all had a common left border corresponding to the 3’ end of IS*801* (position 27,019 of pPsv48C; Fig. 5A), containing the *ori801* where transposition of this element initiates [45]. The right border of the different plasmid derivatives was GAAC (5 clones) or CAAG (8 clones), which were described as consensus tetramers immediately adjacent to insertions of IS*801* and places for one-ended transposition events to finish [22, 46].

The extent and frequency of deletions generated in pPsv48C, both in UPN1007(pRK415) and in UPN1007(pRK3C), was evaluated in clones growing in SNA by a multiplex PCR analysis (Fig. 5B). Additionally, loss of kanamycin resistance indicated the loss of the Km^R^-*sacB* cassette in the largest deletion derivatives (notice that transpositions ending closer from IS*801* result in the deletion of larger DNA fragments from pPsv48C). The 310 suc^R^ clones examined from strain UPN1007(pRK415) retained plasmids of at least 22 kb, all spanning the three TA operons (TA6-8; Fig. 5A). This was expected because the three TA systems are functional in UPN1007 and their loss would predictably result in growth inhibition. However, around half of the clones had lost gene *idi*, indicating the spontaneous loss of this gene in routine culture conditions with a frequency of 0.9 ± 0.3 × 10^−4^. The types of deletions were more varied in the 323 suc^R^ clones of UPN1007(pRK3C), containing functionally inactivated TA systems, with nearly half of the clones losing the RepA replicon and around 80 % (4.4 ± 1.9 × 10^−4^) of them lacking gene *idi* (Fig. 5). Notably, IS*801* was able to transpose the complete length of pPsv48C in both strains (plasmid group F in Fig. 5), although at a low frequency of around 10^−5^, suggesting that IS*801* is capable of mobilizing more than 40 kb of adjacent DNA. Incidentally, the generation of circular deletion variants of pPsv48C mediated by IS*801* also indicates that, as predicted [46], this element transposes by a rolling circle mechanism.

### Toxin-antitoxin systems also contribute to the maintenance of plasmid pPsv48A and to reducing the occurrence of deletions

Because IS*801* is pervasive in *P. syringae* genomes, we wanted to know if deletions mediated by this element also occurred in other plasmids, and whether or not TA systems are contributing to decrease their frequency. For this, we used strain UPN508, a derivative of strain NCPPB 3335 containing plasmid pPsv48A with an insertion of Tn*5*-GDYN1 located at 1.9 kb 3’ of gene *repA* (Fig. 6) [21]. pPsv48A contains only one replicon and Tn*5*-GDYN1 is inserted between two of the five copies of IS*801* in the plasmid, limiting the types and size of deletions that we can detect, although the experimental setting still allowed us to evaluate the possible occurrence of deletions.

**Fig.6.**
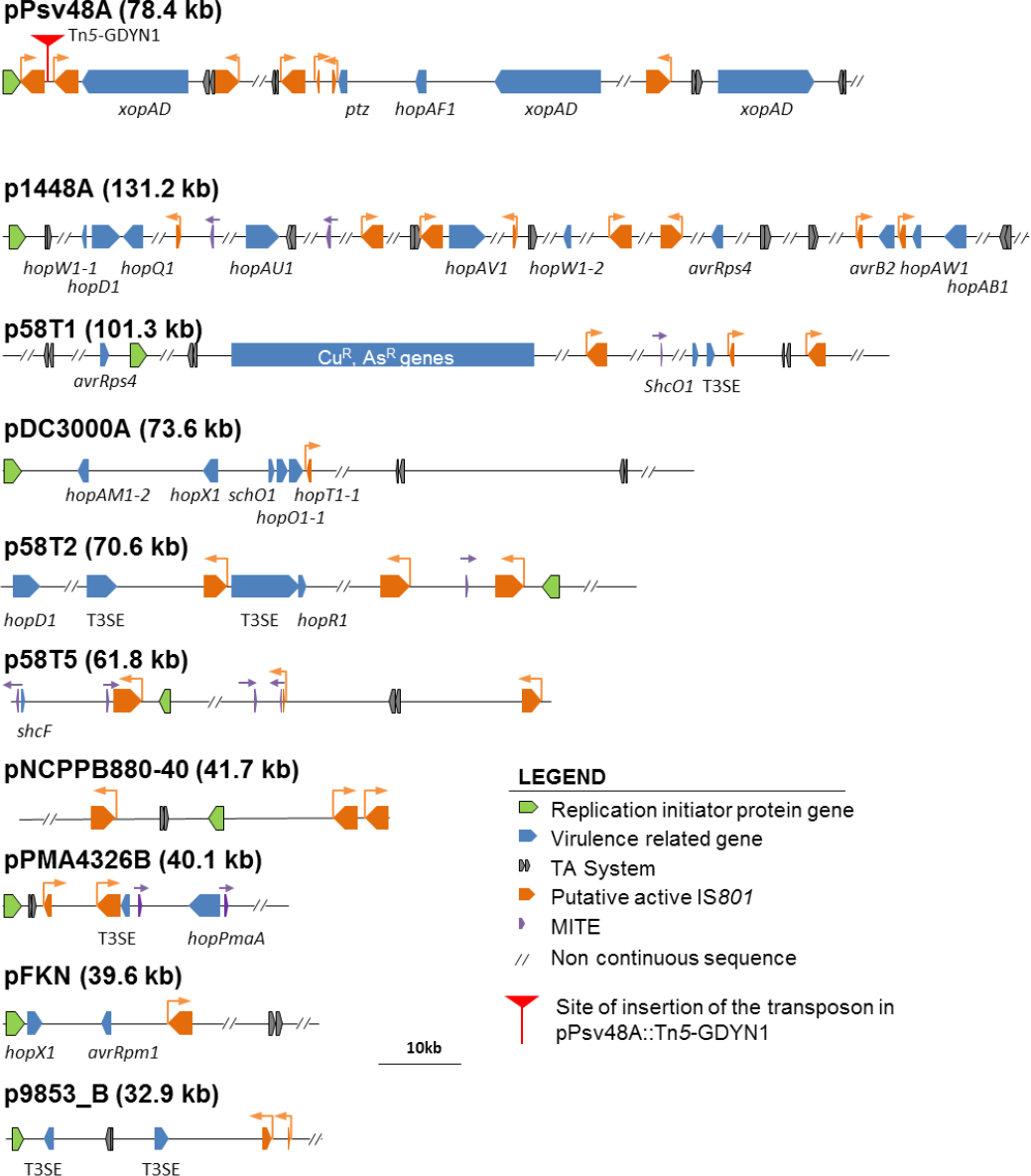
Schematic representation of relevant features found in closed plasmid sequences of *Pseudomonas syringae*. The diagram shows the replicons, virulence genes, TA systems, putative active IS*801* elements and MITES found in closed plasmid fragments are shown.

Strain UPN508(pRK415) generated suc^R^ clones with a frequency of 1.1 ± 0.8 × 10^−4^. From 282 of these suc^R^ clones, plasmid pPsv48A::Tn*5*-GDYN1 was lost in two clones, it contained spontaneous mutations inactivating *sacB* in nine clones, and was reorganized or contained deletions in the remaining ones (Table 3). The majority of the suc^R^ clones, around 90 % of the total, contained derivatives of *ca*. 76 kb; sequencing of three of these clones suggests that they resulted from recombination between the two isoforms of IS*801* flanking the insertion point of Tn*5*-GDYN1 (Table 3), causing its deletion.

**Table 3.**
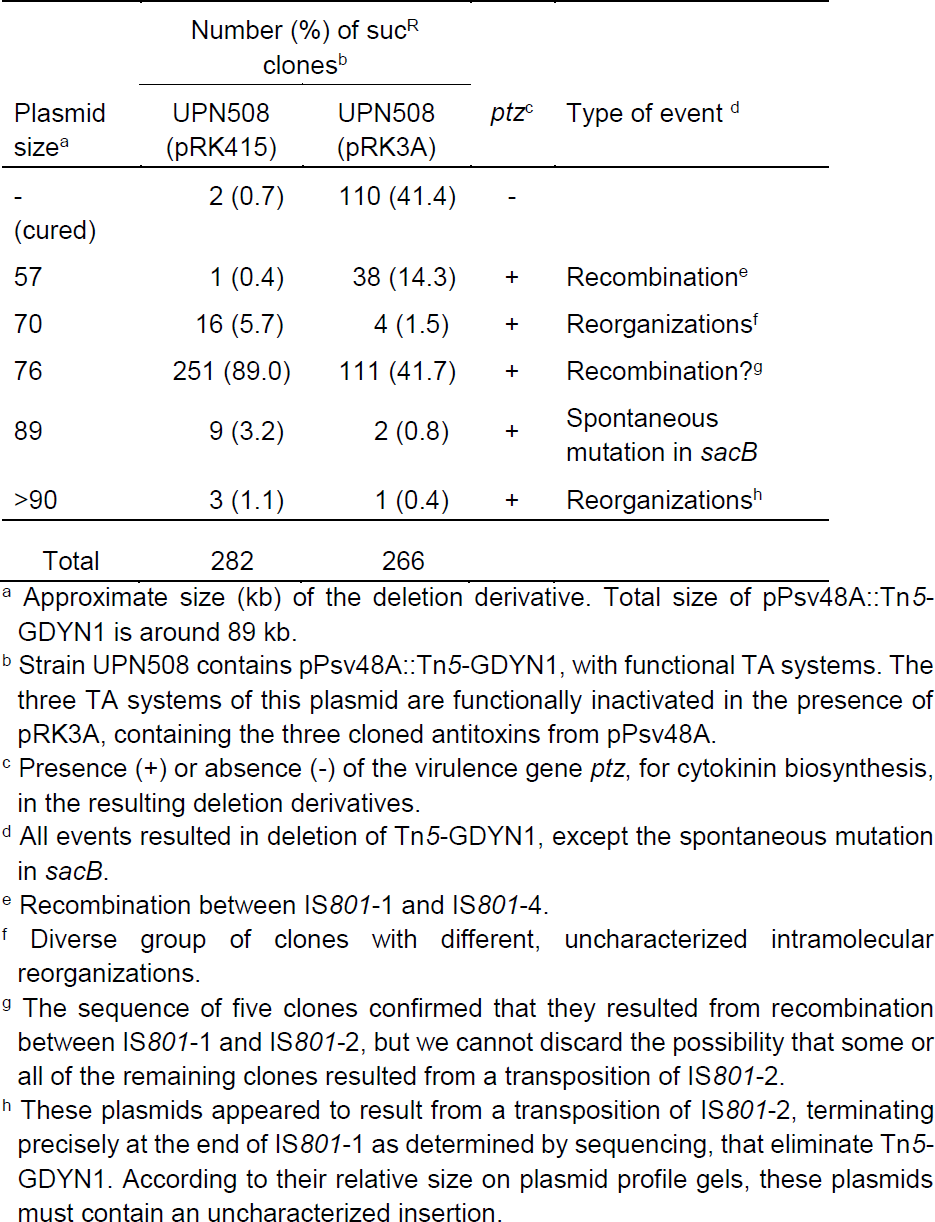
Type and proportion of sucrose-resistant derivatives of pPsv48A::Tn*5*-GDYN1 in the presence or absence of functional toxin-antitoxin systems.

Functional inactivation of the three TA systems, in strain UPN508(pRK3A), lead to only a modest, but significant increase of the frequency of suc^R^ clones to 3.6 ± 1.5 × 10^−4^, and to a dramatic change on the plasmid content of these clones (Table 3). The first major difference was that the frequency of loss of pPsv48A was around 1.5 ± 0.2 × 10^−4^, two orders of magnitude higher than that in UPN508(pRK415) (Table 3). The second major difference was that deletion derivatives of approx. 57 kb, all of which had lost system TA1, appeared around 40 times more frequently than in strain UPN508(pRK415) (Table 3). The frequency of occurrence of the other reorganizations (Table 3) varied no more than four times between both strains. Noticeably, and contrasting with pPsv48C, most of the deletions affecting pPsv48A are likely due to recombination between IS*801* elements instead to one-ended transpositions of IS*801*. This indicates that IS*801* promotes plasmid deletions with high frequency by diverse mechanisms.

### Are *P. syringae* plasmids frequently exposed to IS*801*-mediated deletionsŒ

Many plasmids of *P. syringae* contain virulence genes, and we wondered how often they could be exposed to deletions mediated by IS*801*, and if the presence of this element is associated to the occurrence of TA systems.

We found sequences homologous to IS*801* in 53 out of the 78 available closed plasmid sequences from strains of the *P. syringae* group (including *P. cerasi*; October, 2018), with around two thirds of them containing at least one complete or truncated copy of CRR1. This indicates a frequent occurrence of this mobile element in the *P. syringae* pangenome. The sequence of nine of these plasmids, chosen as examples, contained one to eight copies of *ori801* potentially capable of initiating one-ended transposition (Fig. 6); four of them also contained one to four copies of MITE*Psy1*. Likewise, eight of the nine plasmids harboured at least one putative TA system; an extreme case is p1448A-A (131.2 kb), containing eight *ori801* and seven putative TA systems (Fig. 6). These TA systems are also likely limiting the occurrence of deletions, which could potentially eliminate one or more of the virulence genes included in these plasmids (Fig. 6).

## Discussion

Native plasmids of *P. syringae* and other phytopathogenic bacteria often carry genes contributing to virulence and resistance to bactericides, sometimes being essential for pathogenicity [1, 7, 17, 20, 21, 47, 48]. Although they are generally considered moderately to highly stable in the few tested *P. syringae* strains [21, 30], there is a general lack of knowledge of the molecular mechanisms involved in long-term plasmid stability. Here we show that the virulence native plasmids from *P. syringae* pv. savastanoi NCPPB 3335 use diverse mechanisms to persist in the cell and maintain their physical integrity. Two replicons in pPsv48C [38] confer a high stability level, and particularly RepJ can be maintained, with no apparent plasmid loss, for seven sequential culture transfers in the absence of any identifiable maintenance determinants. Additionally, pPsv48C contains a putative third replicon (RepL), although is apparently non-functional in pseudomonads [21]. Multi-replicon plasmids are common in bacteria, such as the virulence IncF plasmids of clinically relevant *Enterobacteriaceae*, allowing an expansion of their host range [49], but are rarely reported in phytopathogenic bacteria [50]. We also identified three functional TA systems in both pPsv48A and pPsv48C, although they were a major stability determinant only for the first plasmid. The TA systems increased pPsv48A stability by two orders of magnitude and contributed to the maintenance of the virulence gene *ptz* (Table 3), which is essential for the induction of full-size tumours and the development of mature xylem vessels within them [21]. This underscores the different roles played by TA systems and the differential mechanisms used by native plasmids to ensure stable inheritance.

The virulence plasmids pPsv48A and pPsv48C are structurally very fragile, experiencing high frequency intramolecular deletions and reorganizations promoted by the mobile genetic elements MITE*Psy2* and IS*801*. The TA systems carried by these plasmids, however, significantly reduce the accumulation of structural variants by selectively excluding them from the bacterial population. TA systems are bicistronic operons coding for a stable toxin and an unstable antitoxin that neutralises the activity of the toxin [51]. If the operon is lost, for instance due to a deletion, the antitoxin is rapidly degraded and bacterial growth is arrested due to the action of the stable toxin; thus, only cells that did not suffer the deletion and still contain the TA system can grow. Our functional inactivation of the TA systems significantly increased the frequency of the pPsv48C deletions mediated by MITE*Psy2* by 50 times and by three times those mediated by IS*801*. This would indicate that the TA systems might be only moderately successful in preventing deletions mediated by IS*801*. However, we should consider that inactivation of the TA systems lead to a fivefold increase in the loss rate of gene *idi*, which is essential for tumour formation in the plant host [36]. Noticeably, it appears that the loss of gene *idi* was reduced even in those cases where deletion of this gene would not determine loss of any TA system (Fig. 5A). This could be a general feature, because a TA system from a virulence plasmid of *Shigella* spp. favoured the retention of nearby sequences, modulating structural plasmid stability [52]. Likewise, the occurrence of intramolecular deletions and reorganizations of pPsv48A increased three times upon functional inactivation of its TA systems (Table 3); however, this figure is predictably an underestimate because of the limited number and types of events that we could detect with the pPsv48A::Tn*5*-GDYN1 construct used. This phenomenon has been termed post-recombinational killing [52], whereby the occurrence of insertion sequence-mediated rearrangements involving the deletion of TA systems lead to bacterial growth arrest and the consequent exclusion of the reorganized variants from the bacterial population. The occurrence of multiple, apparently redundant, TA systems in plasmids is intriguing. However, plasmids are highly dynamic entities undergoing a continuous trade of genetic material [1, 4]; as such it is feasible that multiple TA systems are selected to ensure the survival of different plasmid fragments. This is clearly exemplified by the 8.3 kb fragment that is “protected” by TA8 (Fig. 3).

In this work, we concentrated on examining the plasmids of strain NCPPB 3335. However, we would expect that the structural fragility of native plasmids and the protective role of TA systems are common phenomena in the *P. syringae* complex, and likely in other plant pathogens, for three main reasons. First, repetitive mobile genetic elements, and particularly IS*801*, are widespread in the *P. syringae* complex, can represent at least one third of diverse native plasmids, and are often associated to virulence genes [21, 22, 25, 30, 53]. IS*801* is remarkable, because it can efficiently transpose with a transposase provided *in trans* and because it follows a rolling circle replicative mechanism, leading to permanent insertions [22, 45, 46]. This implies that any fragment of IS*801* containing *ori801* is potentially mobilizable, that every transposition generates a potentially recombining site, and that one-ended transposition events can immediately lead to the generation of small to very large plasmid deletions. Additionally, other highly repetitive genes, such as the *rulAB* operon for resistance to UV light and many other DNA repair genes, are also commonly associated to virulence and other adaptive genes in *P. syringae* and many other bacteria [54–56]. All these repetitive genetic elements favour the mobility of virulence genes, promoting the high plasticity and adaptability of native plasmids [7, 19–21]; however, at the same time, represent recombination hotspots that can mediate deletion of key virulence genes [57], as highlighted by our results, and of many other adaptive genes. Second, the frequencies of recombination between MITEs and of transposition of IS*801* were very high, suggesting that they could be very active in promoting genomic changes. Nevertheless, and although we observed similar transposition frequencies of IS*801* in two different strains [22], we cannot discount the possibility that these frequencies would be variable among different pathovars or species. Third, and although largely ignored, TA systems are increasingly being found associated to native plasmids in many diverse plant pathogens, including *P. syringae* [20, 58, 59]. It is also noteworthy that most of these plasmids possess several TA systems, as occurs with plasmids from other bacteria (see also Fig. 6) [4, 57, 58].

## Conclusions

*P. syringae* appears to be an opportunistic pathogen whose life cycle primarily occurs in variety of outside-host environments [60]. This sheds uncertainty on the selective forces driving the maintenance of virulence genes, particularly plasmid-borne genes, in the absence of the major selective pressure exerted by the interaction with plant hosts. Indeed, plasmids of *P. syringae* are known to be highly dynamic [1, 7], mostly for the reasons outlined in the previous paragraph. Here we show that TA systems are frequently found in plasmids of *P. syringae* and that they significantly contribute to the maintenance of virulence genes. TA systems have been involved in a disparity of functions including, among others, the stabilization of plasmids and other mobile genetic elements, biofilm formation, modulation of bacterial persistence, resistance to antibacterial compounds, and prevention of large scale deletions in the chromosome, plasmids and episomes [51, 52, 61, 62]. Our results show that genes found in plasmids of the plant pathogen *P. syringae* can be eliminated with high frequency because of plasmid loss and rearrangements mediated by mobile genetic elements. The occurrence of multiple toxin-antitoxin systems in plasmids effectively increase the survival of virulence genes and virulence plasmids in bacterial populations, facilitating their preservation in a diversity of environments lacking the strong selective pressure exerted by the plant host.

## Methods

### Bacterial strains, plasmids and growth conditions

Table 2 summarizes strains, native plasmids and constructions used in this study. LB medium [63] was routinely used for growing both *E. coli* (at 37 ºC) and *Pseudomonas* strains (at 25 ºC). Counter selection of cells carrying the *sacB* gene, which confers lethality in the presence of sucrose, was carried out in nutrient agar medium (Oxoid, Basingstoke, UK) supplemented with 5 % sucrose (medium SNA). When necessary, media were supplemented with (final concentrations, in µg ml^−1^): ampicillin, 100; gentamicin, 12.5; kanamycin, 7 for *P. syringae* and 50 for *E. coli*; tetracycline, 12.5.

### General molecular procedures and bioinformatics

DNA was amplified using a high fidelity enzyme (PrimeStar HS, Takara Bio Inc., Japan), or a standard enzyme (BIOTaq, Bioline, UK), and primers detailed in Table 4. Amplicons were cloned using the CloneJET PCR Cloning Kit (Thermo Scientific) or the pGEM-T Easy Vector System (Promega). Purification of plasmids from *E. coli* was carried out following a boiling method [64] or using a commercial kit (Illustra plasmidPrep Mini Spin Kit, GE Healthcare). For plasmid profile gels, DNA was purified by alkaline lysis and separated by electrophoresis in 0.8 % agarose gels with 1xTAE as described [28]. Plasmids were transferred to *P. syringae* by electroporation [65].

**Table 4.**
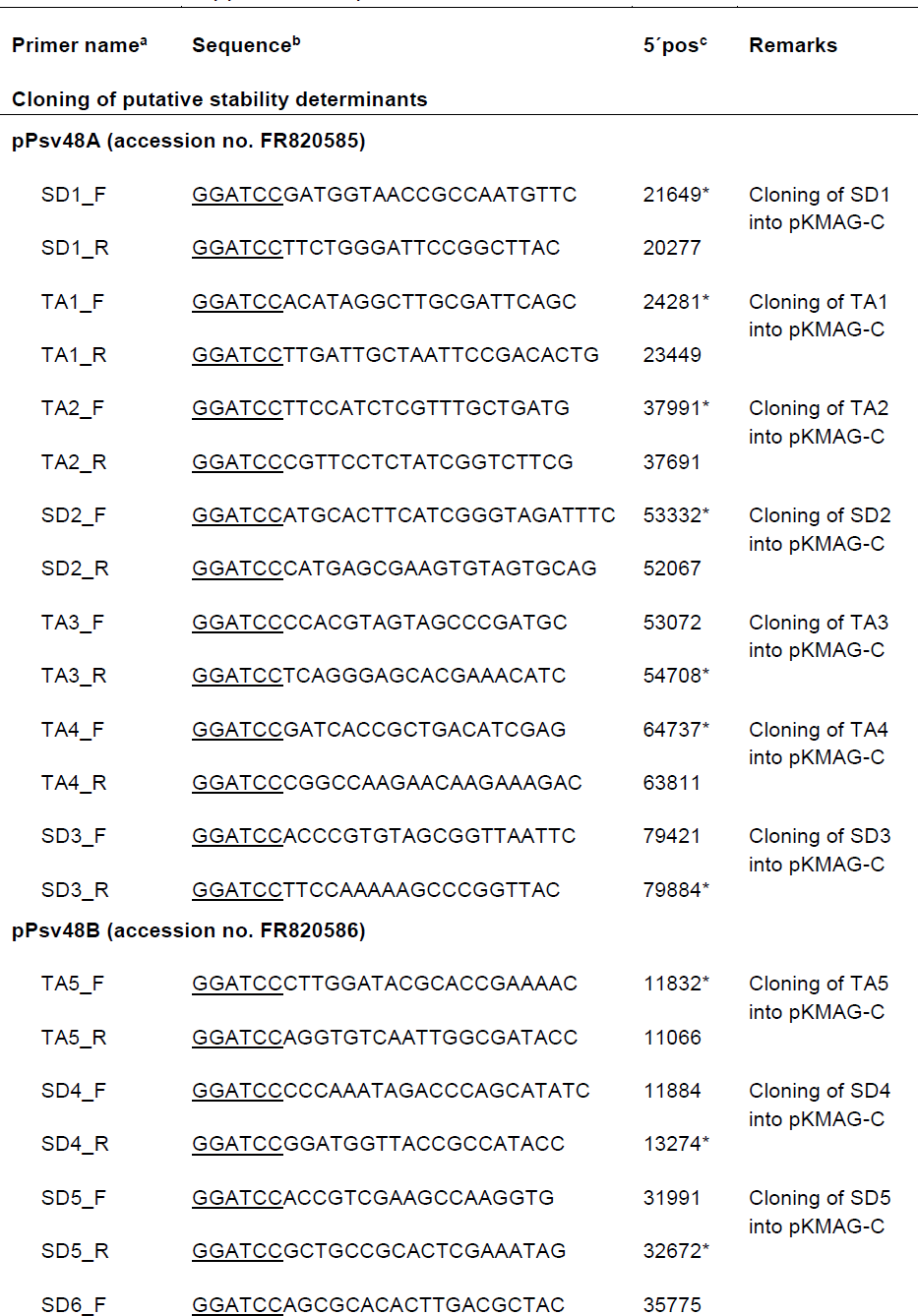
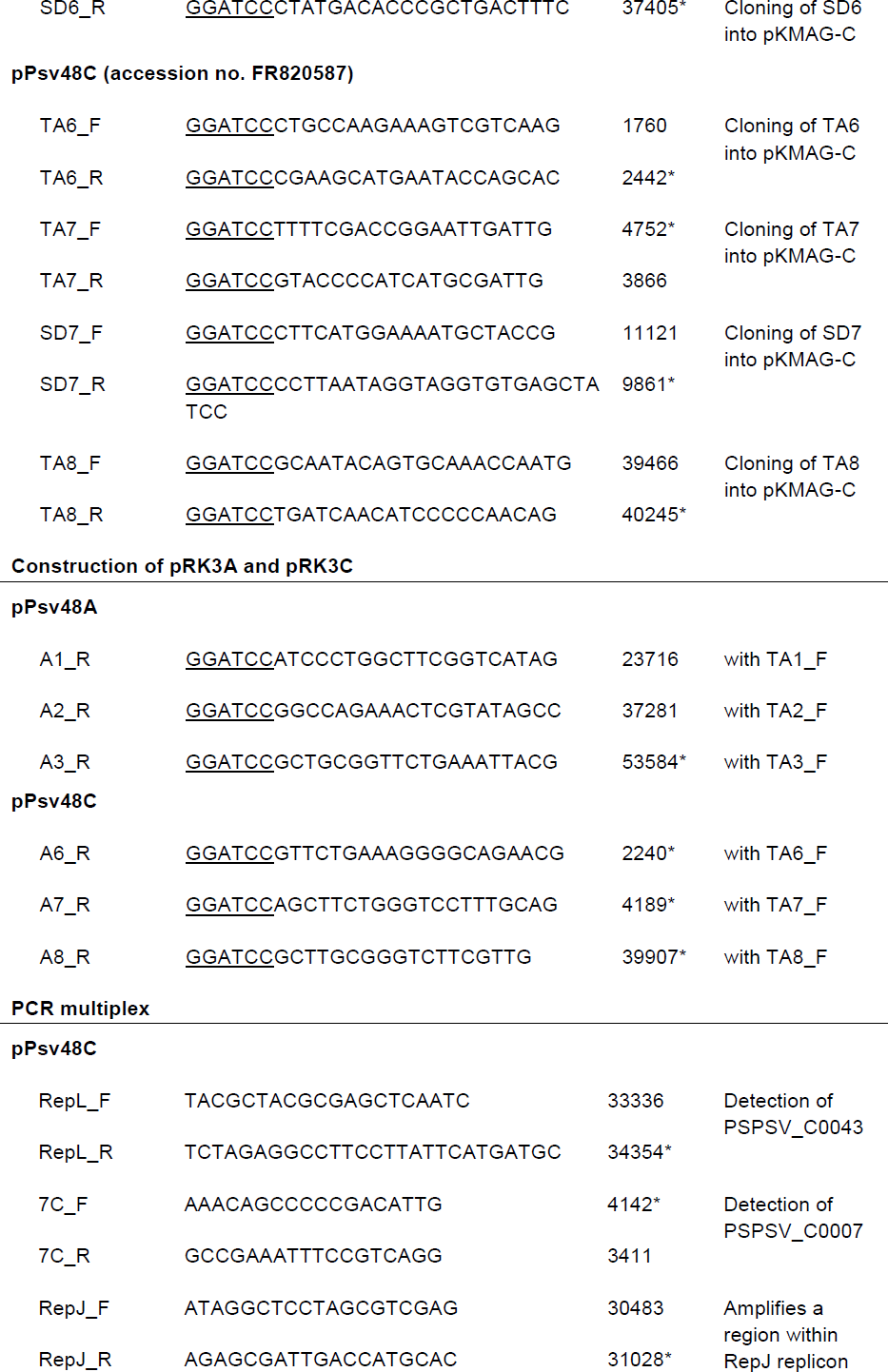
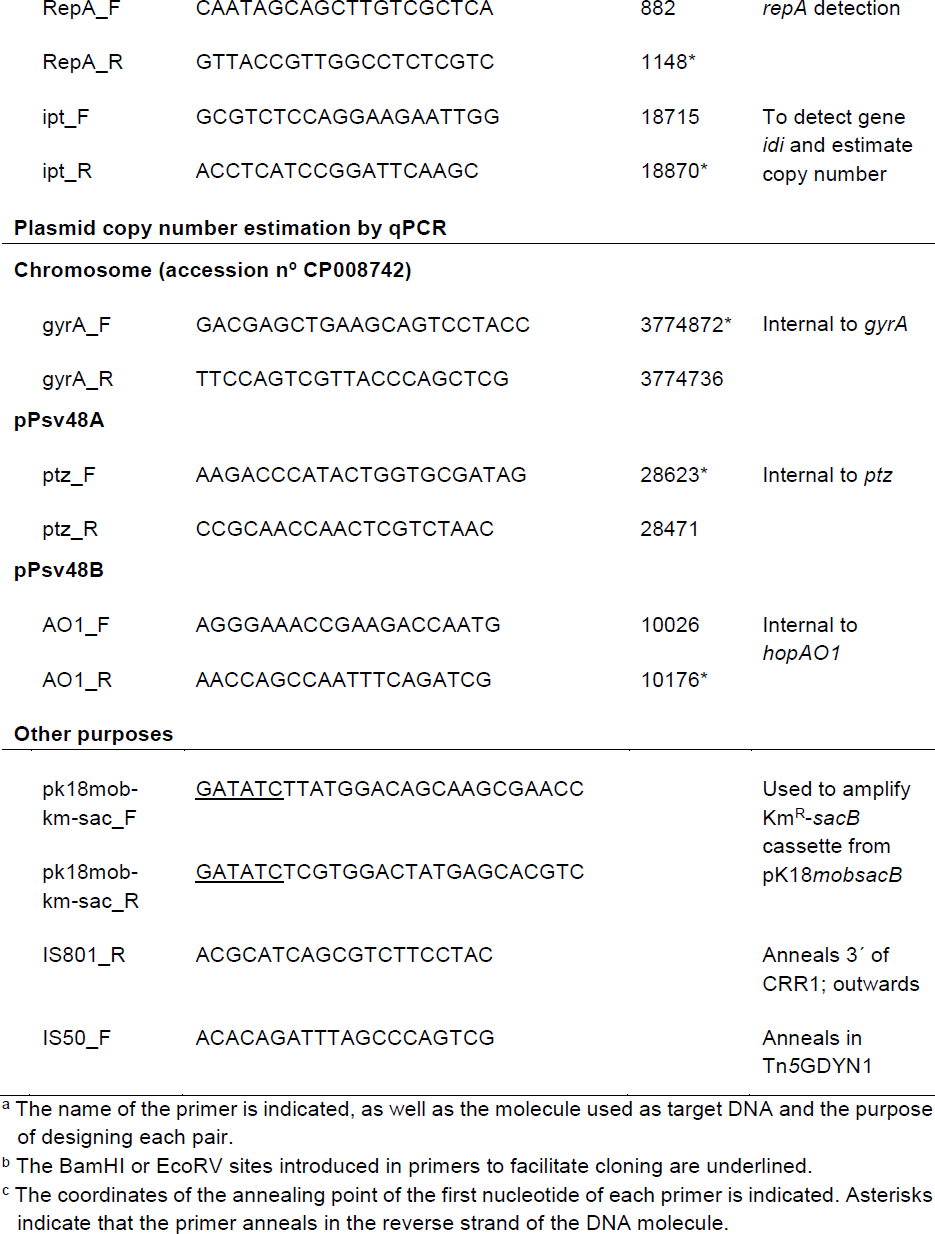
List and application of primers used in this work.

DNA sequences were compared and aligned using the BLAST algorithms [66], as well as the on-line MULTALIN [67] and EMBL-EBI server tools (http://www.ebi.ac.uk/Tools/msa/). The InterPro interface [68] (http://www.ebi.ac.uk/interpro/) was used to search for protein motifs. Nucleotide sequence visualization and manipulation was performed using the Artemis genome browser and ACT [69]. Primers were designed using the Primer3plus software [70].

### Manipulation of native plasmids of *P. syringae* pv. savastanoi

Native plasmids of *P. syringae* pv. *savastanoi* were tagged with Tn*5*-GDYN1 by conjugation using *E. coli* S17.1 as a donor; this transposon carries the levansucrase gene *sacB*, which allows for the identification of derivatives cured of plasmids by selection in medium with sucrose [21, 42]. Sites of Tn*5*-GDYN1 insertion were determined by sequencing of cloned EcoRI fragments containing the Gm^R^ end of the transposon and the adjacent sequences using primer IS50_F (Table 4).

We constructed a derivative of pPsv48C containing a Km^R^-*sacB* cassette, immediately 5’ of the IS*801* isoform (100 nt upstream), as a tool to analyse the diverse deletions generated by the activity of this mobile element. The Km^R^-*sacB* cassette was amplified from pK18*mobsacB* [71] by PCR with specific primers (Table 4), and introduced into an EcoRV site of pPsv48C (position 26,919 in accession no. FR820587) by allelic exchange recombination.

### Estimation of plasmid copy number

Plasmid copy number was estimated by quantitative PCR (qPCR) using as template total DNA purified with the JET flex Genomic DNA Purification Kit (Genomed). qPCR was performed using the CX96^TM^ Real-Time System and analysed using CFX Manager software version 3.0 (BioRad), essentially as described[72]. A ten-fold serial dilution series of DNA was used to construct the standard curve for the single-copy chromosomal gene *gyrA*, used as reference [72], and the plasmids genes *ptz* (PSPSV_A0024;pPsv48A),*hopAO1* (PSPSV_B0010, pPsv48B) and *idi* (PSPSV_C0024, pPsv48C), using the primers indicated in Table 4. Plasmid copy numbers were estimated using the ΔΔCt method [73, 74].

### Identification of putative plasmid stability determinants

For identification of putative stability determinants from plasmids pPsv48A (FR820585), pPsv48B (FR820586) and pPsv48C (FR820587), we manually inspected the annotation of the three plasmids and searched for those CDSs containing terms (stability, partition and related forms), or whose products contained typical domains associated to plasmid maintenance. Additionally, we selected putative toxin-antitoxin operons with a significant score (higher than 70) in the web tool RASTA-bacteria [75]. The complete set of loci identified and tested is summarized in Table 1.

### Replication and stability assays

For functional analyses, the putative stability determinants from the three native plasmids of NCPPB 3335 (Table 1) were amplified by PCR with their own promoters, using specific primers, and cloned as BamHI fragments into the polylinker of vector pKMAG-C (construct 1 in Fig. 2). pKMAG-C replicates in *E. coli* through a p15a replicon and in pseudomonads through the cloned RepA-PFP replicon from pPsv48C [38]. The stability of these constructions, as well as that of the RepA-PFP and RepJ replicons from the pPsv48C plasmid and previously constructed chimeras [38], was tested after transformation into the plasmidless strain *P. syringae* pv. syringae B728a, essentially as described [76]. Briefly, transformants were grown overnight on LB plates with kanamycin, and twenty colonies per clone were collected and resuspended together in 500 µl of Ringer’s solution (1/4 strength; Oxoid, Basingstoke, UK). Serial dilutions were then plated on LB agar to get isolated colonies and, once developed, 100 colonies were picked to LB plates with and without kanamycin to determine the percentage of plasmid-containing colonies (Km^R^). The same procedure was followed to test these constructs in strain UPN912 [36]. The unstable cloning vector pKMAG-C was also included in the analyses as the baseline reference. Experiments were repeated at least six times, with three technical replicates for each of the tested clones. Data were analysed by two-way analysis of variance (ANOVA) followed by Duncan’s multiple range test (P < 0.05) using R Project 3.3.3 (R Core Team (2017); Vienna, Austria).

The stability of the minimal RepJ replicon [38], cloned into pKMAG (construct 4 in Fig. 2), was compared to that of plasmid pPsv48CΔ25, a naturally occurring 5.5 kb deletion derivate of pPsv48C that contains the RepJ replicon plus around 2 kb of downstream DNA, but no other maintenance systems. Both plasmids were maintained in strains derived from NCPPB 3335 and with no other native plasmids. Short-term stability was evaluated as stated above for strain B728a. For long-term stability, three independent LB cultures of each strain were started from single colonies and incubated at 25 ºC with shaking and, after overnight growth, 10 μl of each culture were transferred to 3 ml of LB and incubated in the same conditions. We obtained LB plates containing 200-300 colonies both from the starting culture, immediately after single-colony inoculation, and after seven serial transfers in LB. These colonies were transferred to nylon membranes and analysed by colony hybridization [63], using an internal probe for *repJ*. The number of hybridizing colonies out of the total was scored to assess the prevalence of the RepJ replicon in both populations.

### Inactivation of TA systems and estimation of deletion frequencies

To evaluate the role of TA systems on plasmid maintenance, we proceeded to their functional inactivation, by supplying *in trans* the cognate antitoxins cloned in the broad-host range vector pRK415; resulting in the neutralization of the toxin by the cloned antitoxin, as described [43]. Antitoxin genes PSPSV_A0043, PSPSV_A0032 and PSPSV_A0020 from pPsv48A were amplified by PCR with their own promoters, cloned into pGEM-T Easy, excised as BamHI or NcoI-SacI (for PSPSV_A0032) fragments, and sequentially cloned into the BamHI, NcoI-SacI and BglII sites of vector pME6041, respectively. Primers A1_R and TA3_F were used to amplify these three elements as a single fragment, which was cloned into pJET 2.1 (CloneJET PCR Cloning Kit, Thermo Scientific), excised as a BglII fragment and cloned into the BamHI site of pRK415, downstream of the constitutive P*lac* promoter in the vector, resulting in pRK3A. Essentially the same procedure was followed to clone in tandem and in this order, using primers A6_R and TA8_F, antitoxin genes PSPSV_C0050, PSPSV_C0008 and PSPSV_C0003 from pPsv48C into the vector pRK415, resulting in pRK3C. The integrity and fidelity of all clones was confirmed by nucleotide sequencing.

## Abbreviations

MITE: miniature inverted-repeat transposable element
PFP: pPT23A-family plasmids
SD: Stability determinant
suc^R^: resistant to 5 % sucrose
TA: toxin-antitoxin system

## Declarations

## Acknowledgements

We are indebted to Theresa Osinga for her help with the English language.

## Availability of data and materials

All the data supporting the findings are presented in the manuscript. For raw data, please contact authors for data requests.

## Author’s contributions

LB and JM conceived the study and designed the experiments; LB, MA and ME performed the experiments; LB, CR, and JM analysed the data and interpreted the results; LB and JM drafted the manuscript with contributions from MA and CR; all authors read and approved the final manuscript.

## Ethics approval and consent to participate

Not applicable.

## Consent for publication

Not applicable.

